# Fast protein database as a service with kAAmer

**DOI:** 10.1101/2020.04.01.019984

**Authors:** Maxime Déraspe, Sébastien Boisvert, François Laviolette, Paul H Roy, Jacques Corbeil

## Abstract

Identification of proteins is one of the most computationally intensive steps in genomics studies. It usually relies on aligners that don’t accommodate rich information on proteins and require additional pipelining steps for protein identification. We introduce kAAmer, a protein database engine based on amino-acid k-mers, that supports fast identification of proteins with complementary annotations. Moreover, the databases can be hosted and queried remotely.

## Main

One fundamental task in genomics is the identification and annotation of DNA coding regions that translate into proteins via a genetic code. Protein databases increase in size as new variants, orthologous and paralogous genes are being sequenced. This is particularly true within the microbial world where bacterial proteomes’ diversity follows their rapid evolution. For instance, UniProtKB (Swiss-Prot / TrEMBL) (1) and NCBI RefSeq (2) contain over 100 million bacterial proteins and that number grows rapidly.

Identification of proteins often relies on accurate, but slow, alignment software such as BLAST or hidden Markov model (HMM) profiles (3, 4). Although other approaches (such as DIAMOND (5)) have considerably improved the speed of searching proteins in large datasets, from a database standpoint much can be done to offer a more versatile experience. One such approach would be to expose the database as a permanent service making use of computational resources for increased performance (i.g. memory mapping) and leveraging the cloud for remote analyses via a Web API. Another approach would be to extend the result set with comprehensive information on protein targets to facilitate subsequent genomics and metagenomics analysis pipelines.

Alignment software usually relies on a seed-and-extend pattern using an index (two-way indexing in DIAMOND) to make local alignments between query and target sequences. However, there is a plethora of research techniques to by-pass the computational cost of alignment. Alignment-free sequence analyses usually adopt k-mers (overlapping subsequences of length k) as the main element of quantification. They are extensively used in DNA sequence analyses ranging from genome assemblies (6) to genotyping variants (7), as well as genomics and metagenomics classification (8–10). In the present study, we introduce kAAmer, a fast and comprehensive protein database engine that was named after the usage of amino acid k-mers which differs from the usual nucleic acid k-mers. We demonstrate the usefulness and efficiency of our approach in protein identification from a large dataset and antibiotic resistance gene identification from a pan-resistant bacterial genome.

The database engine of kAAmer is based on log-structured merge-tree (LSM-tree) Key-Value (KV) stores (11). LSM-trees are used in data-intensive operations such as web indexing (12, 13), social networking (14) and online gaming (15, 16). KAAmer uses Badger (17), an efficient implementation in Golang 1 of a WiscKey KV (key-value) store (16). WiscKey’s LSM-tree design is optimized for solid state drives (SSD) and separates keys from values to minimize disk I/O amplification. Disk I/O amplification is typical of LSM-trees due to its vertical design in which keys and values need to be read and rewritten in multiple levels of the tree. Therefore, kAAmer will obtain peak performance with modern hardware such as NVMe 2 SSDs. Furthermore, traditional block devices such as SATA solid-state drives that offer good throughput in input/output (I/O) operations per second (IOPS) will effectively accommodate use cases where many queries are sent simultaneously. A kAAmer database includes three KV stores (see Figure 1A): one to provide the information on proteins (protein store) and two to enable the search functionalities (k-mer store and combination store). The k-mer store contains all the 7-mers found in the sequence dataset and the keys to the combination store, which uniquely serves the combination of proteins held by k-mers. The fixed k-mer size at 7 was chosen to fit on 4 bytes and keep a manageable database size while offering good specificity over protein targets. The k-merized design of a kAAmer database provides an interesting simplicity for the search tasks which will give an exact match count of all 7-mers between a protein query and all targets from a protein database. The result sets using this strategy are not guaranteed to return the same homologous targets that would be obtained with alignment or HMM search and is therefore less suitable for distant homology retrieval (< 50% identity). Nonetheless, kAAmer also supports alignment on the result set without sacrificing speed as shown in Figure 1B. The main drawback of a kAAmer database (in the version at the time of writing: 0.4) is the disk space and time required to build a database that is greater than its benchmarked competitors, although it compares favorably to Ghostz (18) for these parameters.

**Fig. 1.**
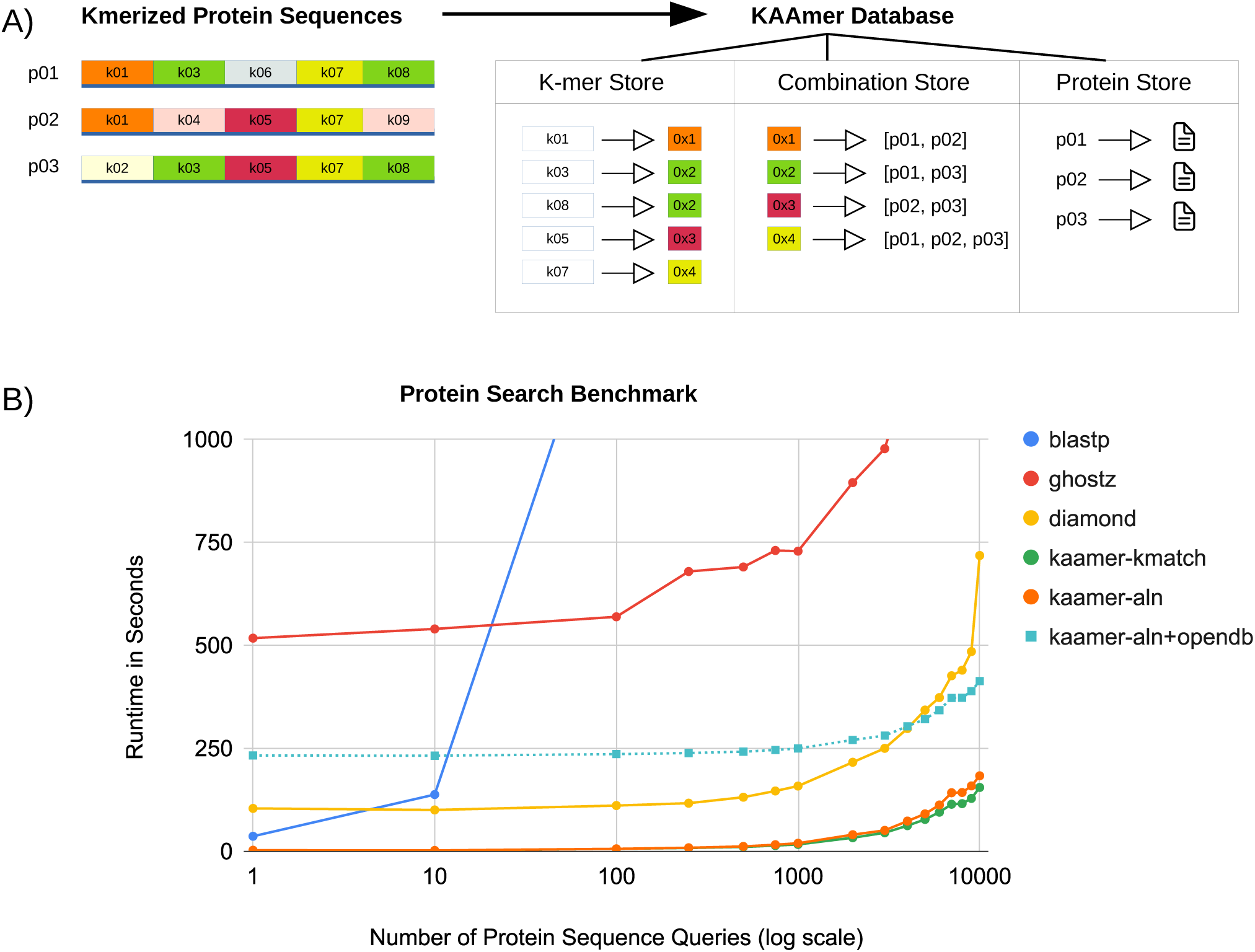
A) Design of a kAAmer database. Three key-value stores are created within a database (K-mer Store, Combination Store, Protein Store). Colours indicate the combination (hash) value that are reused in the combination store. Proteins are numbered (p01, p02, p03) and k-mers are numbered (k01,k02,…,k08). B) Protein search benchmark. Software include blastp (v2.9.0+), ghostz (v1.0.2), diamond (v0.9.25) and kAAmer (v0.4) with and without alignment.

In order to test the efficiency of our database search engine, we used all (114,830,954) the non-fragmented proteins of the UniprotKB (Swiss-Prot / TrEMBL) bacterial proteins dataset (release 2019_08). Sixteen different protein query datasets were randomly and uniquely chosen from the original database, with size ranging from 1 protein to 10,000 proteins. We added the kAAmer search in k-mer match mode (without alignment; named “kaamer-kmatch”) for comparison purposes. We also corrected the kAAmer alignment mode (“kaamer-aln”) in “kaamer-aln+opendb”, by adding the time it took to open the database before running the queries (230 seconds). However, kAAmer’s purpose is to be used as a persistent service so the database opening time becomes insignificant the more you query the database. The four software included in the benchmark are Blastp (v2.9.0+) (3), Ghostz (v1.0.2) (18), Diamond (v0.9.25) (5) and kAAmer (v0.4). Figure 1B illustrates the wallclock times of the alignment software in comparison with kAAmer for protein homology searches. See the Methods section for the hardware used in the benchmarks. We observe that with the larger query datasets (10,000 proteins), kAAmer in alignment mode completes its search and alignments in just 3 minutes 2 seconds (and 2 minutes 26 seconds without alignment). In comparison, Diamond, the second fastest aligner, achieved the 10,000 query task in 11 minutes 57 seconds. Thus, at the maximal benchmarked query size, kAAmer shows an increase in speed of almost 4× (3.93×). When the search incorporates fewer protein queries the gain of kAAmer is more substantial (up to 82× with only 10 protein queries) because Diamond and its double indexing is optimized to perform better when the number of queries increases. It is worth mentioning that when correcting for the database opening (square symbol in Figure 1B.), the kAAmer gain in speed drops and it only surpasses Diamond when there are over 4000 protein queries. However, as stated earlier, kAAmer is rather suited to act as a permanent and flexible database service that will store structured protein information and offer a quick homology search over that protein database. Also, with sufficient random access memory (RAM), data is going to be cached by the operating system (OS) which will increase the performance of kAAmer. For the other benchmarked software, Ghostz took over 33 minutes to realise the task with the 10,000 queries, which is 11 times slower than kAAmer. For Blastp, we stopped the benchmark at 2,000 protein queries since it was already taking over 7 hours to complete the task (at least 700x slower than kAAmer).

In order to accomodate real-use cases we built relevant kAAmer databases and investigated their usage in typical bacterial genomics analyses. It should be noted that annotation of genomes and gene identification rely heavily on the quality of the underlying database. What kAAmer has to offer is the inclusion of the protein information within the database combined with an efficient search functionality to facilitate downstream analyses. Therefore, we also provide utility scripts to demonstrate these use cases. The first use case was to identify antibiotic resistance genes (ARG) in a bacterial genome and test its accuracy related to other ARG finder software. For ARG identification we used the NCBI Bacterial Antimicrobial Resistance Reference Gene Database (v2020-01-06.1) (19) and compared the kAAmer results with the ResFinder (v3.2 and database 2019-10-01) and CARD (v5.1.0) (21) software and database. The query genome is a pan-resistant *Pseudomonas aeruginosa* strain E6130952 (22). Table 1 shows the results of the ARG identification within the query genome by the three software / databases tested. For the majority of antibiotic classes, the results are in agreement between the three databases. Interestingly, three aminoglycoside genes (*aac(6’)-Il, ant(2”)-Ia* and *aacA8*) were only found with kAAmer (NCBI-ARG) and ResFinder. On the other hand, several more antibiotic efflux systems are annotated in CARD and the number of identified efflux proteins in E6130952 goes up to 36 while only 3 were reported by kAAmer (NCBI-ARG) and none by ResFinder. Also 2 genes associated with resistance to peptide antibiotics (arnA, basS) and 2 other (soxR, carA) associated with multiple antibiotic classes were only reported by CARD. Other tested use cases include genome annotation and metagenome profiling as shown in the Methods section.

**Table 1.** Report of the antibiotic resistance genes identification within the pan-resistant Pseudomonas aeruginosa E6130952 strain from kAAmer+NCBI-arg, ResFinder and CARD databases.

| Resistance Gene | Antibiotic Class | kAAmer+NCBI-ARG | ResFinder | CARD |
| --- | --- | --- | --- | --- |
| aac(6')-II | amikacin/kanamycin/tobramycin | 3 | 3 | 0 |
| ant(2'')-Ia | gentamicin/kanamycin/tobramycin | 1 | 1 | 0 |
| aacA8 | aminoglycoside | 1 | 1 (aac(6')-31) | 0 |
| aph(3')-IIb | kanamycin | 1 | 1 | 1 |
| aadA6 | streptomycin | 2 | 2 | 2 |
| blaOXA-2, blaOXA-488 | beta-lactam | 2 | 2 | 2 |
| blaPDC-35 | cephalosporin | 1 | 1 (blaPAO) | 1 (blaPDC-2) |
| fosA | fosfomycin | 1 | 1 | 1 |
| catB7 | chloramphenicol | 1 | 1 | 1 |
| sul1 | sulfonamide | 3 | 3 | 3 |
| mexA, mexE, mexX | efflux | 3 | 0 | 2 (no mexX) |
| other efflux system | efflux | 0 | 0 | 34 |
| arnA, basS | peptide antibiotic | 0 | 0 | 2 |
| soxR, carA | multiple antibiotic class | 0 | 0 | 2 |
| Total | 13 | 19 | 16 | 51 |

In summary, kAAmer introduces a fast and flexible protein database engine to accommodate different genomics analyses use cases. It can be hosted on-premise or in the cloud and be queried remotely via an HTTP API.

## Methods

### Design of kAAmer

KAAmer design was influenced by our requirement that protein databases would be permanently hosted (on premise or in the cloud), queried remotely and would have room to scale as sequence databases grow in size. It also needed to be multithreaded for protein searches and would support alignment for more accurate remote homology findings. We opted for a Key-Value store engine that would reside on disk and be optimized for SSDs. We used the Go programming language for its versatility and efficiency. The Key-Value stores use the Badger (17) engine and protein annotations are encoded using Protocol Buffers (23).

### Database building

KAAmer is first used to build a database in which all amino acid k-mers are associated with proteins in which they are found. It consists of three KV stores to hold the database information (k-mer store, combination store and protein store). The first KV store (k-mer store) keeps the association of every k-mer (key) with a hash value (key length: 8 bytes) that is the entry to the combination store. The k-mer size is fixed at 7 amino acids to fit k-mer keys onto 32 bits (4 bytes) and thus maintain a manageable final database size while keeping a k-mer size long enough for specificity. The second KV store (combination store) is used to hold all the unique sets of protein identifiers. The method used to build this store can relate to the flyweight design pattern, the hash consign technique, and the coloured de Bruijn graphs (7, 8). Indeed, hash values are reused to access identical objects and therefore minimize memory usage. The set of protein identifiers are the keys to the last store (protein store) which contains the protein information found in the raw annotation file. The raw input file can be either in the EMBL format, GenBank format, TSV format or in FASTA format.

### Querying a database

Once we have a database, we expose it with the kAAmer server that listens over HTTP for incoming requests. The benefits of using such a service are two fold. First, the database is opened once and is memory mapped to increase the performance of protein searches. Second, the kAAmer server can be hosted virtually anywhere, in the cloud for instance, and be queried remotely by the kAAmer client. Note that it is preferable that the latency (time required for a message to be transported over HTTP) between the server and client be as low as possible. KAAmer supports protein query and translated DNA query from FASTA input as well as short reads sequences (like Illumina) in FASTQ format.

### Benchmark protein alignment software

To build the benchmark on the UniProtKB bacterial proteins database, we randomly and uniquely extracted multiple sets of sequences, with the number of sequences ranging from 1 to 10 thousand. Each set of sequences was in its own FASTA file to be queried with the different alignment software included in the benchmark. The benchmark for all four software (Blastp (v2.9.0+), Ghostz (v1.0.2), Diamond (v0.9.25) and kAAmer (v0.4)) was run on nodes geared with 32 cores (Intel(R) Xeon(R) CPU E5-2667), 120 GB of RAM and with a SATA III connected SSD. The maximum number of results for each query was set to 10 and no threshold was provided. All software were run with default parameters, except for the number of threads set to 32 and the maximum number of results per query at 10.

### Other kAAmer use cases

Apart from the antibiotic resistance gene (ARG) identification use case, we also provide two demonstrations of kAAmer usage in bacterial genome annotation and metagenome profiling. The use cases are documented at https://github.com/zorino/kaamer_analyses and a Python script is provided for each one of the analyses. For the genome annotation, we used the chromosomal sequence of the same *Pseudomonas aeruginosa* strain (E6130952) as in the antibiotic resistance genes identification. The kAAmer database that was used for the homology detection is a subset of RefSeq from the *Pseudomonadaceae* family which is available from the kAAmer repository (see Data availability) along with other Bacterial family databases. Essentially the genome annotation script parses the kAAmer results and produces a GFF (General Feature Format) annotation file giving some threshold on the protein homology. The other use case is the profiling of a metagenome based on the MGnify database of the human gut (24). MGnify includes protein annotations from gene ontology, enzyme commission and kegg pathways, among others. The metagenome profiling script will parse the results and produce a summary file by annotation that counts the presence and abundance of each feature.

### Data availability

We have built a repository where one can download prebuilt kAAmer database versions of common protein datasets useful in bacterial genomics and metagenomics. The repository is available at https://kaamer.genome.ulaval.ca/kaamer-repo/ and includes datasets from the NCBI, the EBI and other popular data sources.

### Code availability

The code is available at https://github.com/zorino/kaamer under the Apache 2 license and the documentation can be found at https://zorino.github.io/kaamer/.

## Acknowledgements

This study was financed by the Canada Research Chair in medical genomics (JC). MD was supported by the Fonds de recherche du Québec - Santé (#32279). The authors thank Juan-Manuel Dominguez and Charles Burdet for their comments. Computations were performed under the auspices of Calcul Québec and Compute Canada. The operations of Compute Canada are funded by the Canada Foundation for Innovation (CFI), the National Science and Engineering Research Council (NSERC), NanoQuébec and the Fonds Québécois de Recherche sur la Nature et les Technologies (FQRNT).

## Declaration

The authors declare no competing interests.

## Footnotes

1 Go programming language (https://golang.org/)

2 Non-Volatile Memory Express

## Bibliography

1. The Uniprot Consortium. UniProt: a worldwide hub of protein knowledge. Nucleic Acids Research, 47(D1):D506–D515, jan 2019. ISSN 0305-1048. doi: 10.1093/nar/gky1049.

2. Nuala A O’Leary, Mathew W Wright, J Rodney Brister, Stacy Ciufo, Diana Haddad, Rich McVeigh, Bhanu Rajput, Barbara Robbertse, Brian Smith-White, Danso Ako-Adjei, Alexander Astashyn, Azat Badretdin, Yiming Bao, Olga Blinkova, Vyacheslav Brover, Vyacheslav Chetvernin, Jinna Choi, Eric Cox, Olga Ermolaeva, Catherine M Farrell, Tamara Goldfarb, Tripti Gupta, Daniel Haft, Eneida Hatcher, Wratko Hlavina, Vinita S Joardar, Vamsi K Kodali, Wenjun Li, Donna Maglott, Patrick Masterson, Kelly M McGarvey, Michael R Murphy, Kathleen O’Neill, Shashikant Pujar, Sanjida H Rangwala, Daniel Rausch, Lillian D Riddick, Conrad Schoch, Andrei Shkeda, Susan S Storz, Hanzhen Sun, Francoise Thibaud-Nissen, Igor Tolstoy, Raymond E Tully, Anjana R Vatsan, Craig Wallin, David Webb, Wendy Wu, Melissa J Landrum, Avi Kimchi, Tatiana Tatusova, Michael DiCuccio, Paul Kitts, Terence D Murphy, and Kim D Pruitt. Reference sequence (RefSeq) database at NCBI: current status, taxonomic expansion, and functional annotation. Nucleic acids research, 44(D1):D733–45, jan 2016. ISSN 1362-4962. doi: 10.1093/nar/gkv1189.

3. Christiam Camacho, George Coulouris, Vahram Avagyan, Ning Ma, Jason Papadopoulos, Kevin Bealer, and Thomas L Madden. BLAST+: architecture and applications. BMC bioinformatics, 10:421, dec 2009. ISSN 1471-2105. doi: 10.1186/1471-2105-10-421.

4. S R Eddy. Profile hidden Markov models. Bioinformatics (Oxford, England), 14(9):755–63, 1998. ISSN 1367-4803. doi: 10.1093/bioinformatics/14.9.755.

5. Benjamin Buchfink, Chao Xie, and Daniel H Huson. Fast and sensitive protein alignment using DIAMOND. Nature methods, 12(1):59–60, jan 2015. ISSN 1548-7105. doi: 10.1038/nmeth.3176.

6. P A Pevzner, H Tang, and M S Waterman. An Eulerian path approach to DNA fragment assembly. Proceedings of the National Academy of Sciences of the United States of America, 98(17):9748–53, aug 2001. ISSN 0027-8424. doi: 10.1073/pnas.171285098.

7. Zamin Iqbal, Mario Caccamo, Isaac Turner, Paul Flicek, and Gil McVean. De novo assembly and genotyping of variants using colored de Bruijn graphs. Nature Genetics, 44(2):226–232, feb 2012. ISSN 1061-4036. doi: 10.1038/ng.1028.

8. Sébastien Boisvert, Frédéric Raymond, Élénie Godzaridis, François Laviolette, and Jacques Corbeil. Ray Meta: scalable de novo metagenome assembly and profiling. Genome Biology, 13(12):R122, 2012. ISSN 1465-6906. doi: 10.1186/gb-2012-13-12-r122.

9. Derrick E Wood and Steven L Salzberg. Kraken: ultrafast metagenomic sequence classification using exact alignments. Genome Biology, 15(3):R46, 2014. ISSN 1465-6906. doi: 10.1186/gb-2014-15-3-r46.

10. Brian D. Ondov, Todd J. Treangen, Páll Melsted, Adam B. Mallonee, Nicholas H. Bergman, Sergey Koren, and Adam M. Phillippy. Mash: Fast genome and metagenome distance estimation using MinHash. Genome Biology, 17(1):132, 2016. ISSN 1474760X. doi: 10.1186/s13059-016-0997-x.

11. Patrick O’Neil, Edward Cheng, Dieter Gawlick, and Elizabeth O’Neil. The log-structured merge-tree (LSM-tree). Acta Informatica, 33(4):351–385, jun 1996. ISSN 0001-5903. doi: 10.1007/s002360050048.

12. Fay Chang, Jeffrey Dean, Sanjay Ghemawat, Wilson C. Hsieh, Deborah A. Wallach, Mike Burrows, Tushar Chandra, Andrew Fikes, and Robert E. Gruber. Bigtable. ACM Transactions on Computer Systems, 26(2):1–26, jun 2008. ISSN 07342071. doi: 10.1145/1365815.1365816.

13. Sanjay Ghemawat and Jeffrey Dean. LevelDB, 2011.

14. Facebook. RocksDB, 2013.

15. Biplob Debnath, Sudipta Sengupta, and Jin Li. SkimpyStash. In Proceedings of the 2011 international conference on Management of data - SIGMOD ‘11, page 25, New York, New York, USA, 2011. ACM Press. ISBN 9781450306614. doi: 10.1145/1989323.1989327.

16. Lanyue Lu, Thanumalayan Sankaranarayana Pillai, Andrea C Arpaci-Dusseau, and Remzi H Arpaci-Dusseau. WiscKey: Separating Keys from Values in SSD-conscious Storage. In 14th {USENIX} Conference on File and Storage Technologies ({FAST} 16), pages 133–148, Santa Clara, CA, feb 2016. {USENIX} Association. ISBN 978-1-931971-28-7.

17. Dgraph Labs. Badger, 2017.

18. Hongwei Ge, Liang Sun, and Jinghong Yu. Fast batch searching for protein homology based on compression and clustering. BMC bioinformatics, 18(1):508, nov 2017. ISSN 1471-2105. doi: 10.1186/s12859-017-1938-8.

19. Michael Feldgarden, Vyacheslav Brover, Daniel H Haft, Arjun B Prasad, Douglas J Slotta, Igor Tolstoy, Gregory H Tyson, Shaohua Zhao, Chih-Hao Hsu, Patrick F McDermott, Daniel A Tadesse, Cesar Morales, Mustafa Simmons, Glenn Tillman, Jamie Wasilenko, Jason P Folster, and William Klimke. Validating the AMRFinder Tool and Resistance Gene Database by Using Antimicrobial Resistance Genotype-Phenotype Correlations in a Collection of Isolates. Antimicrobial agents and chemotherapy, 63(11), nov 2019. ISSN 1098-6596. doi: 10.1128/AAC.00483-19.

20. Ea Zankari, Henrik Hasman, Salvatore Cosentino, Martin Vestergaard, Simon Rasmussen, Ole Lund, Frank M Aarestrup, and Mette Voldby Larsen. Identification of acquired antimicrobial resistance genes. The Journal of antimicrobial chemotherapy, 67(11):2640–4, nov 2012. ISSN 1460-2091. doi: 10.1093/jac/dks261.

21. Brian P Alcock, Amogelang R Raphenya, Tammy T Y Lau, Kara K Tsang, Mégane Bouchard, Arman Edalatmand, William Huynh, Anna-Lisa V Nguyen, Annie A Cheng, Sihan Liu, Sally Y Min, Anatoly Miroshnichenko, Hiu-Ki Tran, Rafik E Werfalli, Jalees A Nasir, Martins Oloni, David J Speicher, Alexandra Florescu, Bhavya Singh, Mateusz Faltyn, Anastasia Hernandez-Koutoucheva, Arjun N Sharma, Emily Bordeleau, Andrew C Pawlowski, Haley L Zubyk, Damion Dooley, Emma Griffiths, Finlay Maguire, Geoff L Winsor, Robert G Beiko, Fiona S L Brinkman, William W L Hsiao, Gary V Domselaar, and Andrew G McArthur. CARD 2020: antibiotic resistome surveillance with the comprehensive antibiotic resistance database. Nucleic Acids Research, oct 2019. ISSN 0305-1048. doi: 10.1093/nar/gkz935.

22. Jianhui Xiong, Maxime Déraspe, Naeem Iqbal, Sigmund Krajden, William Chapman, Ken Dewar, and Paul H. Roy. Complete Genome of a Panresistant Pseudomonas aeruginosa Strain, Isolated from a Patient with Respiratory Failure in a Canadian Community Hospital. Genome Announcements, 5(22), jun 2017. ISSN 2169-8287. doi: 10.1128/genomeA.00458-17.

23. Google. Protocol Buffers, 2008.

24. Alex L Mitchell, Alexandre Almeida, Martin Beracochea, Miguel Boland, Josephine Bur- gin, Guy Cochrane, Michael R Crusoe, Varsha Kale, Simon C Potter, Lorna J Richardson, Ekaterina Sakharova, Maxim Scheremetjew, Anton Korobeynikov, Alex Shlemov, Olga Kun- yavskaya, Alla Lapidus, and Robert D Finn. MGnify: the microbiome analysis resource in 2020. Nucleic Acids Research, nov 2019. ISSN 0305-1048. doi: 10.1093/nar/gkz1035.

